# Comprehensive regulatory networks for tomato organ development based on the genome and RNAome of microTom tomato

**DOI:** 10.1101/2022.12.01.518646

**Authors:** Jia-Yu Xue, Hai-Yun Fan, Zhen Zeng, Yu-Han Zhou, Shuai-Ya Hu, Sai-Xi Li, Ying-Juan Cheng, Xiang-Ru Meng, Fei Chen, Zhu-Qing Shao, Yves Van de Peer

## Abstract

MicroTom tomato has a short growth cycle and high transformation efficiency, and is a prospective model plant for studying organ development, metabolism, and plant-microbe interactions. Here, with a newly assembled reference genome for this tomato cultivar and abundant RNA-seq data derived from tissues of different organs/developmental stages/treatments, we constructed multiple gene co-expression networks, which will provide valuable clues for the identification of important genes involved in diverse regulatory pathways during plant growth, e.g., arbuscular mycorrhizal symbiosis and fruit development. Additionally, non-coding RNAs, including miRNAs, lncRNAs and circRNAs were also identified, together with their potential targets. Interacting networks between different types of non-coding RNAs (miRNA-lncRNA), and non-coding RNAs and genes (miRNA-mRNA and lncRNA-mRNA) were constructed as well. Our results and data will provide valuable information for the study of organ differentiation and development of this important fruit. Lastly, we established a database (http://eplant.njau.edu.cn/microTomBase/) with genomic and transcriptomic data, as well as details of gene co-expression and interacting networks on microTom, and this database should be of great value to those who wants to adopt microTom as a model plant for research.

## Introduction

Tomato (*Solanum lycopersicum*) is one of the most popular fruits (although usually referred to as a vegetable) in the world. In 2020, its annual worldwide production was estimated at about 186 millions of tons and has been increasing every year (http://www.fao.org/faostat). Tomato is also an emerging model plant system for developmental biology and, in some cases, can be an better option than *Arabidopsis thaliana*, e.g., for studies of fruit development ^1, 2^, metabolism ^3–5^, plant-pathogen interactions ^6, 7^, and arbuscular mycorrhizal (AM) symbiosis ^8, 9^.

MicroTom is a tomato cultivar and currently a widely applied experimental model plant for lab studies. This cultivar has a smaller size than regular tomato cultivars (e.g., Heinz 1706 and M82), and together with a shorter growth cycle and higher transformation efficiency makes it one of the best choices for a lab model among tomato cultivars. However, a high quality genome has been lacking for this model cultivar, despite the fact that hundreds of tomato cultivars/accessions have been (re-)sequenced already ^4,10–12^. Nowadays, developmental biologists rely on the Heinz tomato genome as a reference, whereas functional experiments are mainly performed using microTom as transformation system. Many re-sequencing studies have indicated considerable sequence diversity among cultivar/accession genomes ^4, 12–14^ and the newly developed pan-genome strategy revealed the existence of specific genes only belonging to certain cultivars/accessions ^11, 15, 16^. Such observations suggest that the different genetic background between the reference genome and others might complicate experiments, for instance in cloning target genes, potentially leading to failure of the entire experimental design. Therefore, the availability of a high quality microTom genome was badly needed.

In this study, we provide a high-quality genome of microTom and conducted comparative genomic analysis with the previously published ‘Heinz’ tomato genome. Additionally, together with large amounts of RNA-seq data sequenced by this study and collected from public databases, we present the RNAome landscape of microTom across different organ/developmental stages/treatments and performed comprehensive analyses of the transcriptome of microTom protein-coding genes, with respect to (aspects of) gene expression and alternative splicing (AS). With the reference genome and abundant gene expression data, we constructed co-expression networks for tomato organ development, which will provide valuable information for the regulation of organ development in the life cycle. Non-coding RNAs were also identified by combining different sequence strategies, and their interaction networks were predicted. Finally, we constructed a database (microTomBase, http://eplant.njau.edu.cn/microTomBase), available for data download, online search, and demonstration of analytical results. This database will be invaluable for researchers who employ the microTom tomato as the lab model plant.

## Results

### Assembly of the microTom genome and comparative genomics between microTom and Heinz

Using a combination of 92.40 Gb Nanopore data and 54.38 Gb Illumina data, we first obtained a microTom genome assembly of 799 Mb with 60 contigs (contig N50 = 41.37 Mb). Then, using the genome of Heinz as a reference, 58 microTom contigs were further merged to 12 corresponding pseudo-chromosomes. Protein-coding gene annotation of the microTom assembly captures 98.57 % of the embryophyta BUSCO (odb10) genes, with 97.89 % single copy genes and 0.68% of duplicates. These results indicate that our assembled microTom genome has reached a high standard of quality and completeness (**Table 1**).

**Table 1.** Statistics and comparison of microTom and Heinz genomes.

|  | MicroTom |  |  |  | Heinz 1706 (SL4.0) |  |  |  |
| --- | --- | --- | --- | --- | --- | --- | --- | --- |
|  | Contigs |  | Pseudomolecules |  | Contigs |  | Pseudomolecules |  |
|  | Size(bp) | Number | Size(bp) | Number | Size(bp) | Number | Size(bp) | Number |
| N50 | 41,367,923 | 8 | 66,788,879 | 6 | 6,007,830 | 37 | 65,269,487 | 6 |
| Maximum length | 68,578,620 |  | 94,893,795 |  | 26,291,688 |  | 90,863,682 |  |
| Total number | 60 |  | 12 |  | 448 |  | 12 |  |
| Gaps number | 47 |  |  |  | 435 |  |  |  |
| Assembled length | 798,935,606 |  |  |  | 782,475,302 |  |  |  |
| BUSCO completeness | 98.57% |  |  |  | 97.40% |  |  |  |
| Gene models number | 35,213 |  |  |  | 34,690 |  |  |  |

With the assistance of three full-length transcriptome RNA-seq, 12 ribo-minus RNA-seq data, and 69 poly-A enriched RNA-seq datasets derived from different organs/tissues under different developmental stages and/or treatments (See Methods and Materials) from this and previous studies ^3, 17^, 35,213 protein-coding genes were annotated in microTom. Among those, 31,891 genes (90.57%) received transcriptomic data support from at least one RNA-seq sample. Although microTom has a similar number of protein-coding genes than Heinz’s (34,690), these two cultivars share only 2/3 one-to-one orthologous genes according to synteny analysis and bi-directional blast (**Figure 1a**), yet each cultivar encodes thousands of specific genes, comprising real specific genes, paralogs by duplications, and genes missed by annotation (mainly in Heinz probably due to inadequate RNA-seq data) (**Supplementary Figure S1**). For the two cultivars, 4,449 microTom genes were “hiding” in the Heinz genome (gene length coverage and DNA sequence identity > 90%), only not annotated; while the corresponding number of Heinz genes not annotated in microTom dropped to a half. This comparison indicates that adequate and diverse transcriptomic data are critical to a more comprehensive genome annotation.

**Figure 1.**
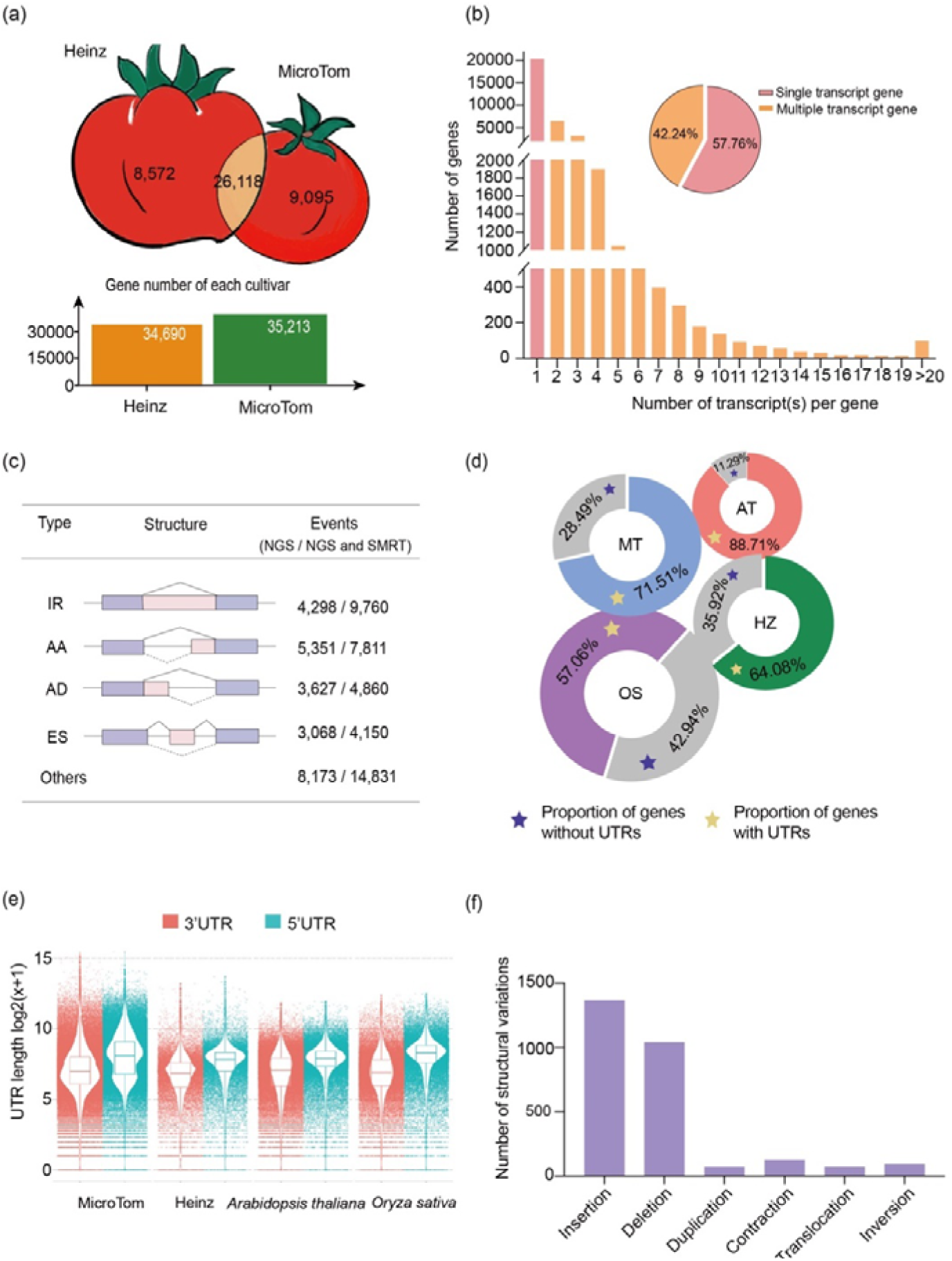
Genome annotation of microTom and comparison with Heinz and other species. **(a)** Protein-coding gene numbers for Heinz and microTom varieties. Shared one-to-one orthologs in microTom and Heinz genomes are presented in the cross section of the two tomato cartoon pictures. **(b)** Proportion of genes with multiple transcripts and genes with different transcript numbers. **(c)** Categories and frequency of AS events in microTom. IR, intron retention; AA, alternative adaptor; AD, alternative donor; ES, exon skipping. **(d)** Proportion of genes with annotated UTRs in microTom (MT), Heinz (HZ), *Arabidopsis thaliana* (AT), an d*Oryza sativa* (OS). **(e)** Dot plot of annotated UTR lengths in microTom, Heinz, *Arabidopsis thaliana* and *Oryza sativa*. **(f)** Detected structural variation events of microTom, compared with Heinz. The minimum size of insertion, deletion, duplication and contraction is 150 bp, the minimum size of inversion is 1 Kb, and the minimum size of translocation is 10 Kb.

While alternative splicing (AS) has been widely detected in eukaryotes to generate functionally divergent transcripts from a common parental gene ^18^, the Heinz genome did not provide multi-transcript annotation for protein-coding genes. By incorporating dozens of RNA-seq datasets from PacBio and Illumina platforms, 14,873 (42.24%) microTom protein-coding genes were identified to have multiple transcripts (**Figure 1b**), with an average of two transcripts per gene. Among all AS transcripts, intron retention accounts for the largest proportion (23.57%), followed by alternative acceptor (18.86%), and alternative donor (11.74%) (**Figure 1c**). The expression patterns of all transcripts were examined based on the transcriptomic data (**Supplementary Table S1**). Among the AS transcripts, the majority (> 70%) showed obvious differential expression in different organs and/or developmental stages (**Supplementary Table S1**), suggesting that these AS transcripts may be functionally distinct and regulate organ differentiation and development in a more prevalent manner than we previously thought.

Due to the large amount of transcriptomic data, more comprehensive information for the untranslated regions (UTRs) was also available. About 71.28% of microTom genes have annotated UTRs, showing an obviously higher percentage than that in Heinz and the rice and genomes (**Figure 1d**), and the medium lengths of 5’ and 3’ UTRs are 125 bp and 271 bp, respectively (**Figure 1e**). The annotation of UTRs should provide useful information for gene regulation and expression studies.

Comparing the genomes of microTom and Heinz, over one million single nucleotide polymorphism (SNP) sites and 40 thousand insertion-deletions (indels) (< 150 bp) were detected. Additionally, a great number of larger structural variations were also identified (**Figure 1f**), such as bigger-sized genomic insertions, deletions, duplications, contractions, translocations, and inversions. These small and large structural variations indicate genomic divergence between the two tomato cultivars, which may affect experimental results that demand high accuracy, e.g. SNPs and indels in the coding regions would greatly affect the design of CRISPR guide RNAs.

### Co-expression networks

Co-expression networks underlying the development of organs and response to stimuli were inferred using transcriptome data derived from 22 different tissues/organs and at different developmental stages and/or different treatments (three replicates for each sample). All expressed genes were classified into 123 modules (**Figure 2a**, **Supplementary Figure S2** and **Table S2**), whereby each module contains genes that share a similar expression pattern, and possibly function in the same regulatory network, defining and/or regulating a specific phenotype. When a specific phenotype is defined, the corresponding module can be identified, and the hub genes, as well as the entire network linked to the phenotype can be extracted. This way, unknown genes in the pathway can be identified, potentially providing important clues of targets and gene interactions.

**Figure 2.**
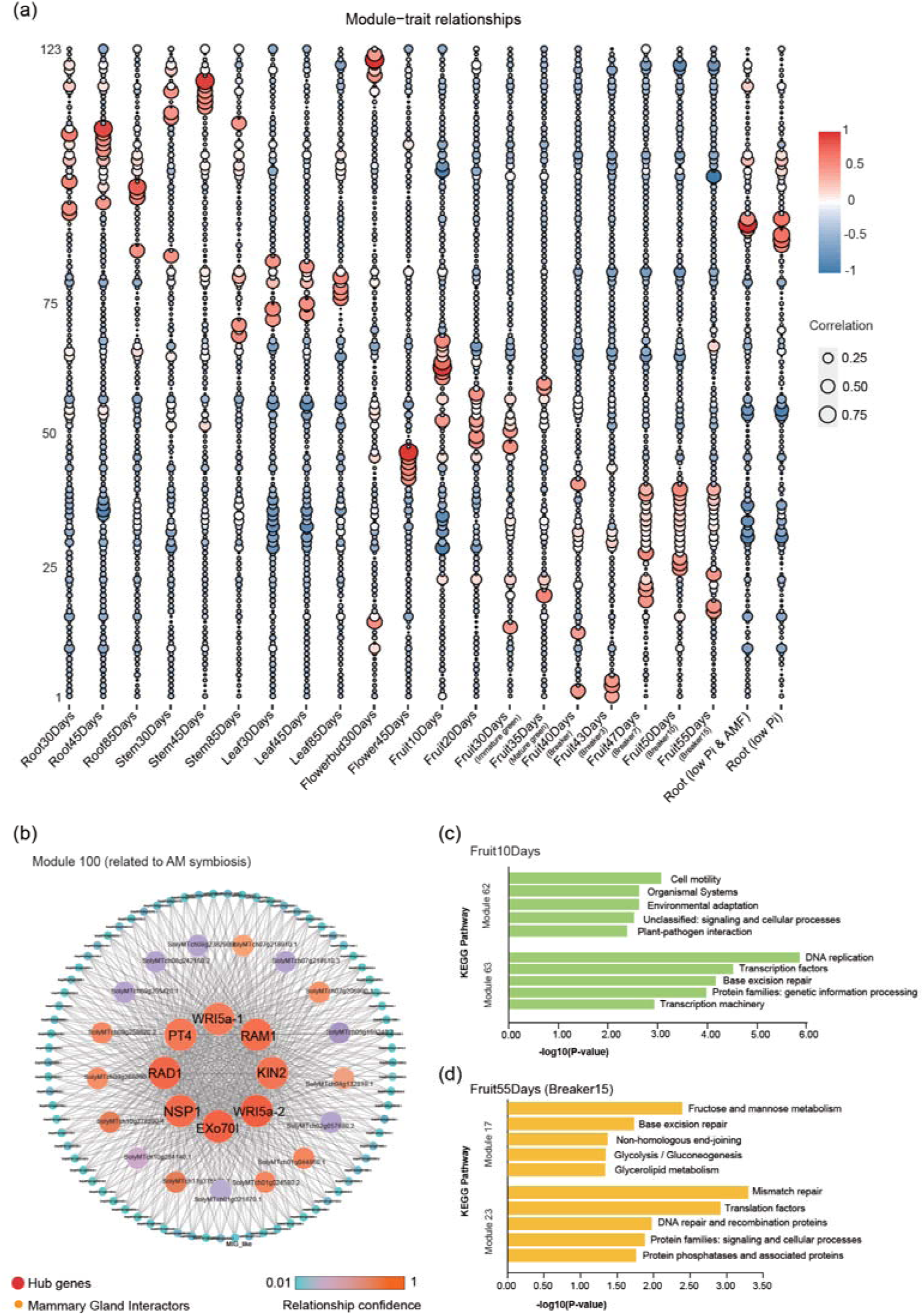
MicroTom co-expression networks established based on 22 transcriptome samples of different organs/developmental stages/treatments. **(a)** 123 modules and their correlation with 22 phenotypes or molecular processes. **(b)** Module 100 and its network, significantly correlated with AM symbiosis. Gene names (instead of IDs) indicate that these genes have been functionally characterized and involved in the AM symbiosis. Threads indicate correlation between genes. **(c)** Top five enriched KEGG pathways in the early stage of fruit development (10 days after anthesis, “Fruit10Days” on Figure 2a) in the most correlated Module 62 and 63.**(d)** Top five enriched KEGG pathways in the fully mature stage of fruit development (55 days after anthesis, “Fruit55Days (Breaker15) on Figure 2a) in the most correlated Module 17 and 23.

For instance, our results indicate that the AM symbiosis in microTom (Root, low Pi & AMF) is significantly associated with Module 100 (**Figure 2a** a n d**2b**, **Supplementary Table S2**). It could be noticed that several important genes of known functions in the AM symbiosis pathway are present, and the majority of them are even hub genes in the network, including PT4, NSP1, RAD1, RAM1, Exo70I, KIN2, and two WRI5a orthologous (designated as WRI5a-1 and WRI5a-2). It is well-known that RAM1, RAD1 and NSP1 are regulators relatively upstream in the AM symbiosis pathway ^19–23^, and WRI5a is even considered as a master regulator ^24^, so they should have broad associations with other genes in the same pathway. Therefore, this co-expression result makes good sense, not only in identifying the right (expected) genes, but likely also in suggesting other critical, potentially even hub, genes. In this case, other unknown genes in the pathway can also be identified through such expression association (guilt by association), and co-expression analysis should serve as an effective means to explore unknown functional genes in pathways.

Tomato is one of the, if not the most important model plant for fruit development studies. Therefore, we collected transcriptomes derived from fruits of nine different developing stages (10, 20, 30, 35, 40, 43, 47, 50, 55 days after anthesis and three replicates each), and observed fruits at different developing stages showing specific associations with different transcriptional modules (**Figure 2a**). For instance, fruits of 10 days after anthesis are associated with modules 62 and 63 (**Figure 2a** and **Supplementary Table S2**), the genes of which are functionally enriched in the categories of “Cell motility”, “Organismal Systems”, “DNA replication” and “Transcription factors”, suggesting that early development of tomato fruits mainly involves cell division, cell proliferation, and regulatory network development of this organ (**Figure 2c**); whereas fruits of 55 days after anthesis are associated with modules 17 and 23 (**Figure 2a** and **Supplementary Table S2**), the genes of which are functionally enriched in the categories “Fructose and mannose metabolism”, “Base excision repair”, “Mismatch repair” and “Translation factors”, suggesting biosynthesis of metabolites, which mainly seems to occur in fully mature fruits (**Figure 2d**). These different modules indicate distinct major regulatory priorities at different developmental stages of tomato fruits.

### Diverse non-coding RNAs, their targets and interactions

Non-coding RNAs, including miRNA, lncRNA and circRNA, were comprehensively annotated using multiple RNA-seq data sets. These datasets were generated from six development stages/organs of microTom tomato plants (root, stem, leaf, flower, green fruit and red fruit), and under phosphorus deficiency and AM symbiosis status. A total of 210 miRNAs were identified from the sRNA datasets, of which 164 belong to 48 known tomato miRNA families in miRBase (**Figure 3a**). We identified 19 tomato miRNA families previously documented in the miRBase with increased family sizes, with one to seven more members discovered for each family. Notably, five miRNA families that have not been documented in tomato (only documented in other plants) were identified in this study, namely miR157, miR1446, miR1886, miR2111 and miR3627. Additionally, 46 novel miRNAs were identified (**Figure 3a**). The lengths of microTom miRNAs range from 20-nt to 24-nt, with 21-nt miRNAs occupying the largest proportion. Expression analysis revealed that most of the identified miRNAs were organ-/developmental stage-specific or were only expressed upon stimulation by Pi deficiency and/or AM symbiosis. Only a few of them could be detected in multiple samples, e.g. miR403-3p, miR9472-5p and miR166a/g.

**Figure 3.**
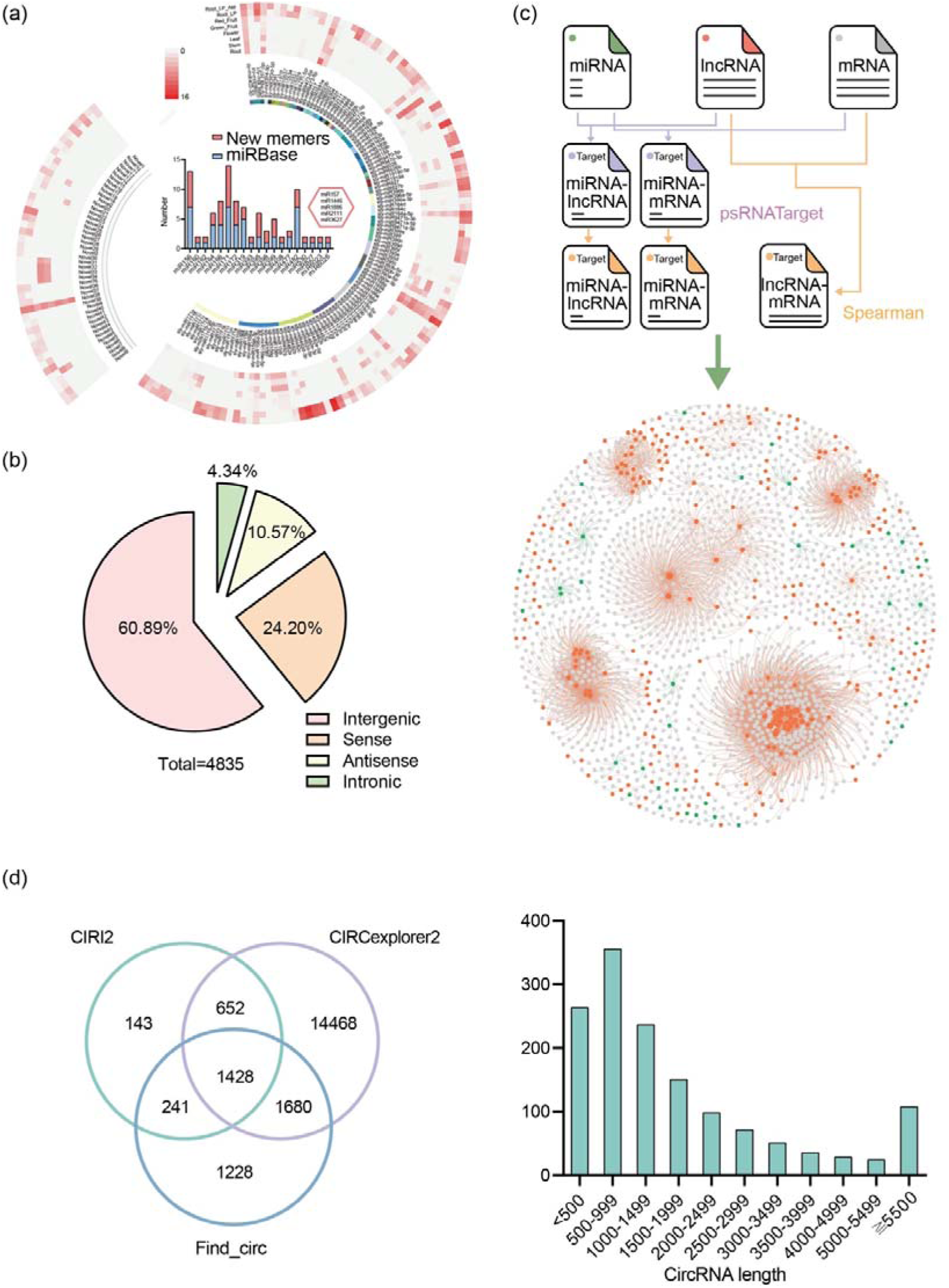
Non-coding RNAs and interaction networks in microTom genome. **(a)** Identification of the microTom miRNAs. The right panel represents known miRNA families and their members identified in this study and their expression, and the left panel represents novel miRNA families. The circle heatmap shows miRNA expression (outer to inner): 1. in roots symbiotic with AM fungi for six weeks under low phosphorus condition, 2. roots without AM fungi under low phosphorus condition, 3. red fruits, 4. green fruits, 5. flowers, 6. leaves, 7. stems, and 8. roots under normal phosphorus condition. The histogram represents the new members identified in the known miRNA families in microTom, the red part is the newly discovered members, and the blue part indicates the existing members of miRBase. The hexagon represents miRNA families previously not identified in tomatoes but identified in other plants. **(b)** Proportion of four different types of lncRNAs. **(c)** An integrated interaction network between miRNAs, lncRNAs and mRNAs. The purple line in the flow chart represents miRNA target gene prediction using psTargetscan, and the orange line represents spearman analysis. The green, red and gray dots represent miRNAs, lncRNAs and mRNAs, respectively. **(d)** The Venn diagram represents the comparison of circRNAs predicted using three software. **(e)** Distribution of circRNA lengths.

Altogether, 4,835 lncRNAs were annotated from microTom transcripts, and over 2,944 of them (60.89%) were transcribed from intergenic regions, whereas 720, 511 and 210 lncRNAs, were transcribed from the genomic regions overlapping with protein-coding genes at the sense/antisense strand, or from the intronic sequences, respectively (**Figure 3b**). The length of microTom lncRNAs ranges from 200 to 15,073 nt, with an average length of 986 nt. Similar to miRNAs, the majority of lncRNAs showed development stages/tissue-specific or stimulus induced expression. To explore the potential regulatory roles of miRNAs and lncRNAs, pair-wise interaction analyses were performed for combinations of miRNA-mRNA, miRNA-lncRNA and lncRNA-mRNA. The results showed that 266 mRNAs and seven lncRNAs could be predicted as targets of miRNAs with a significant negative expression correlation. Significant expression correlation was also observed for 8,672 lncRNA-mRNA pairs, suggesting potential regulatory relationships. Putting these predicted interactions together, networks integrating miRNAs, lncRNAs, and mRNAs were constructed (**Figure 3c** and **Supplementary Table S3**), and these networks should serve as a basic resource for elucidating the regulatory roles of miRNAs and lncRNAs in tomato development and response to stimuli.

A comprehensive annotation of circRNA was performed by incorporating the sequencing results from this study and our previous study ^17^. A total of 19,840 circRNAs, supported by three independent software tools were identified by mapping the sequencing reads onto the microTom genome. The lengths of identified 1,428 high-confidence circRNAs ranged from 197 to 27,820 nt, with an average of 1,428 nt.

### Construction of microTom genomic and RNA-omic database

To make our microTom genomic data and analytical results conveniently accessible, we constructed a database (microTomBase, http://eplant.njau.edu.cn/microTomBase), providing online search and download possibilities of our data and results (**Figure 4**). Through gene ID or sequence blast searches, interesting genes can be found, together with detailed information, including genomic positions, gene sequence with detailed structure information (UTRs and CDSs), functional annotation, multi-transcripts and expressional profiles. Information of non-coding RNAs and their targets as well as interacting networks is also provided under the ‘non-coding RNA’ section, which can also be accessed through blast search. All microTom protein-coding genes are ascribed to 123 co-expression modules and can be found under the “Co-expression” section. Genome assemblies and annotation files of the microTom tomato are available for download. The microTom database will facilitate comparative genomics, transcriptional regulation and functional studies for tomatoes.

**Figure 4.**
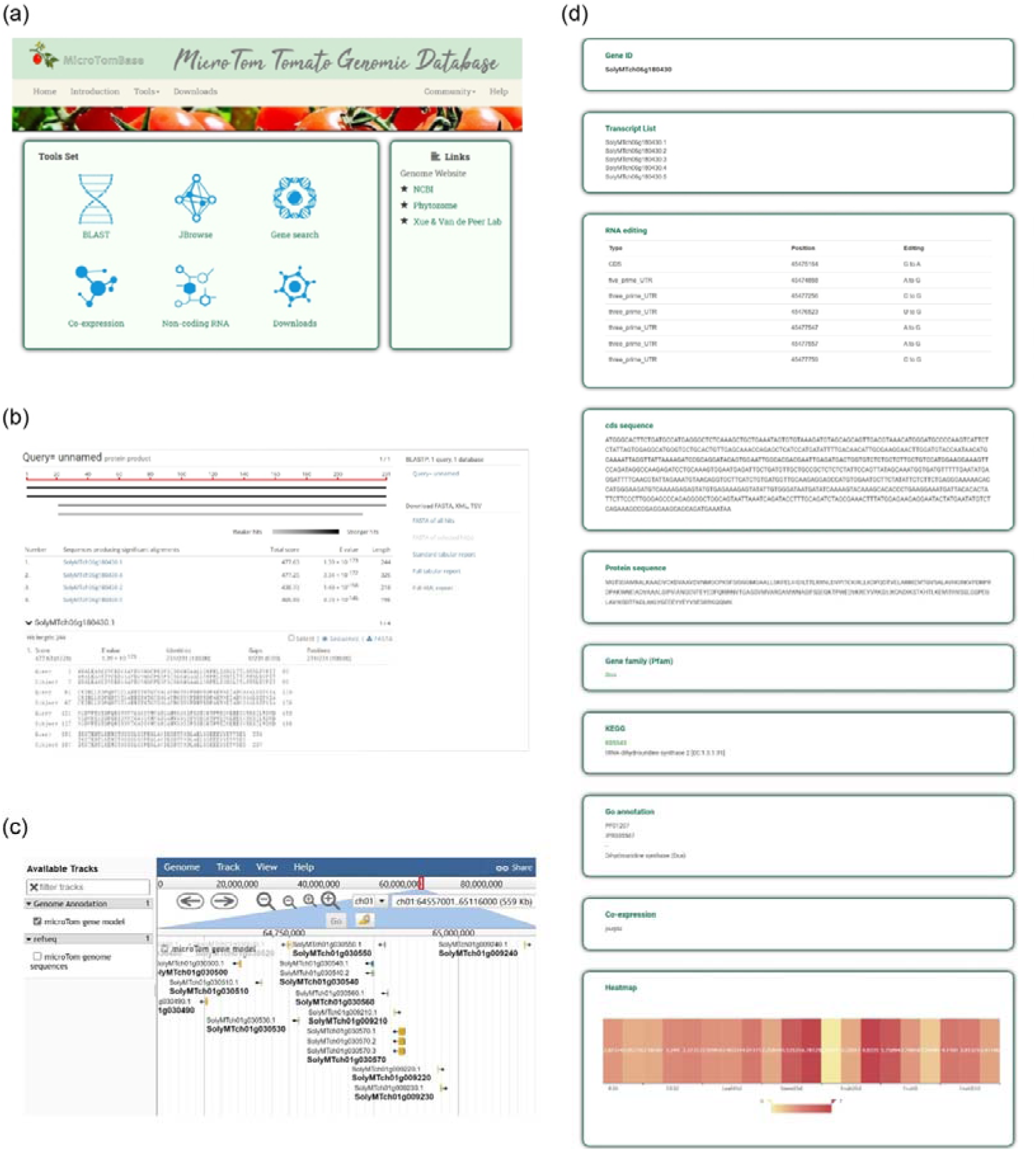
Demonstration of the database MicroTomBase. (**a**) Homepage of MicroTomBase and the main functions, providing service/information of sequence blast, genomic position anchoring, gene ID search, co-expression modules, non-coding RNAs and data download. **(b)** The BLAST tool of MicroTomBase. Users can input genomic, PEP, or CDS sequences as the query sequence and the resulting alignment scores are ranked from high to low. **(c)** The JBrowse tool for visualization of microTom genomic details, including gene visualization interface and detailed data on individual genes. **(d)** Detailed information about genes, including protein and CDS sequence, RNA editing sites, KEGG and GO annotation, gene family, expression profile and co-expression module.

## Discussion

Tomato has been playing an increasingly important role as a model plant for studies of fruit development, plant disease resistance, and symbiosis. Despite being the most employed cultivar, and larges amounts and wide diversities of multi-omic data being available, microTom tomato until now lacked a high quality genome. This study fills this gap by providing a high quality microTom genome assembly, obtained by a combination of Nanopore and Illumina sequencing. Moreover, we integrated a diversity of transcriptome data (obtained from both public resources and our own data) to achieve a comprehensive annotation and profound analyses for this genome, reporting more genes, more and longer UTRs, multiple AS transcripts, co-expression networks, non-coding RNAs and their interaction networks, results which should all serve as a great resource for further genetic and functional studies.

AS is considered as a mechanism to generate multiple transcripts with different functions ^18^, and a number of studies have reported its extensiveness in organismal life cycles, e.g. organ differentiation and development ^25, 26^, disease ontogenesis ^27, 28^ and response to abiotic and biotic stimuli ^29–31^. However, the Heinz tomato genome did not provide information for AS. The microTom genome annotation fills this gap by including multiple transcripts generated by AS. Over 40% of microTom genes are alternatively spliced and the majority of AS transcripts shows differential expression under different conditions, suggesting a broad functional role of AS.

Other than the assist at in-depth genome annotation, the large amounts of transcriptomes also provide data for our co-expression network analyses, which classified microTom transcripts into different expressional modules, and accordingly established diverse networks associated with different phenotypes (different organs, developmental stages and treatments). Potential genes with important functions that have not been recognized can thus be identified based on their association and applied for further functional characterization. Therefore, our co-expression networks should provide useful information for a more precise targeting of candidate genes related to specific phenotypes in the tomato life cycle.

Non-coding RNAs are extensively involved in the post-transcriptional regulation of gene expression through different mechanisms, and play important roles in a variety of life processes on plant growth and development, stress resistance, and interaction with pathogenic or beneficial microbes ^32, 33^. In tomato, high-throughput sequencing of miRNAs of microTom have been reported ^17, 34–36^, and functional roles of miRNAs in fruit developmental regulation have been characterized, e.g., miRNA156 ^37^ and miRNA159 ^38^. The profiles of lncRNAs and circRNAs of microTom have also been investigated, but by only a few studies ^17, 39^. Moreover, these studies merely focused on non-coding RNAs at certain specific developmental stages or treatments, while a full picture of tomato non-coding RNAs is still lacking. In this study, we comprehensively annotated miRNAs, lncRNAs and circRNAs in the microTom genome by integrating multiple datasets generated from diverse organs, developmental stages and treatments (**Supplementary Table S3**), and our results greatly extended the non-coding RNA list of tomato. Furthermore, the expression profile of non-coding RNAs among different samples indicate that most of the identified miRNAs and lncRNAs were specifically expressed in different organ-and developmental stage or under AM symbiosis, suggesting a conditional induction of these non-coding RNAs. Furthermore, non-coding RNAs can regulate the expression of protein-coding genes, directly or coordinatively affecting the translation and homeostasis of mRNAs ^40^. By constructing the interaction networks of the microTom miRNAs, lncRNAs and mRNAs, we uncovered highly complicated regulatory relationships between non-coding RNAs and their target mRNAs, which provides novel insights into the post-transcriptional regulation of protein-coding genes in microTom.

In summary, this study presents a high-quality genome for the microTom tomato, and a comprehensive annotation for both coding and non-coding genes, including splice variants and UTRs. Taking advantage of large amounts of transcriptomes, the gene expression profile and the co-expression networks are constructed, which will provide clues for the identification of novel genes involved in diverse pathways. Non-coding RNAs, including miRNAs, lncRNAs and circRNAs, as well as their potential targets, were identified and interacting networks were constructed, which will provide valuable information for future studies dedicated to exploring the post-transcriptional regulation of plant development by non-coding RNAs. All these findings suggest diverse and complicated modifications and regulations at the RNA level. All these results have been integrated into our online resource, which we hope, could provide an invaluable resource for researchers studying tomatoes and employing tomato as a model plant.

## Materials and methods

### Plant material and data sources

Seedlings of the *Solanum lycopersicum* (cv. ‘microTom’) were grown in plastic pots filled with a mixture of sterilized sand/gravel (1:1 ratio) in a climate-controlled growth room with 16 h light at 24°C and 8 h dark at 22°C. Roots, stems, leaves, flower, immature green fruits, and mature red fruits were collected from plants grown in nutrient-rich soil for six weeks, and used for RNAome sequencing. Green and red fruits were classified by color chart “USDA Visual Aid TM-L-1” (USDA, Agricultural Marketing Service, 1975).

We also downloaded 75 Illumina RNA-seq datasets and two PacBio RNA-seq datasets involving different tissues/organs and different developmental stages and/or different treatments from our previous study ^17^, and a study performed by others ^3^. Twelve microTom tomato proteomic datasets were retrieved from the PRIDE Archive database (Proteomics identifications database, https://www.ebi.ac.uk/pride/archive/).

### Library construction and sequencing

Fresh leaves of four-week old microTom that were grown in half-strength Murashige and Skoog basal medium with sucrose and phytagel were collected, and the genomic DNA was sequenced on the Nanopore platform using a MinION R9 flow cell, and the Illumina NovaSeq6000 platform with an insert size of 450bp at Benagen (Wuhan, China).

Fresh tissues of roots, stems, leaves, flower, immature green fruits and mature red fruits of plants grown in nutrient-rich soil were subjected to total RNA extraction using TRIzol (Invitrogen, Carlsbad, USA) according to the manufacturer’s protocol, which were used for subsequent RNA sequencing using different strategies. Samples from each tissue were subjected to construct a ribo-minus RNA library and small RNA library for RNA sequencing, whereas a mixture of the six samples were used to construction a PacBio SMRT library and a circular RNA library. The ribo-minus RNA libraries, small RNA libraries and circular RNA library were sequenced using the Illumina HiSeq platform at Novogene Co., Ltd. (Tianjin, China), and the PacBio SMRT library was sequenced using the PacBio Sequel System at Novogene Co., Ltd. (Tianjin, China).

### Genome assembly and assessment

Firstly, the raw Nanopore reads were corrected and assembled into contigs by NextDenovo-v2.5.0 (https://github.com/Nextomics/NextDenovo) with default parameters. After that, Nanopore reads were further mapped into primary contigs by software Minimap2-v2.17 ^41^ with the parameter ‘-ax map-ont’, followed by Nextpolish-v1.4.1 ^42^ to polish contigs. Meanwhile, the Illumina paired-end reads were processed to remove adaptor and low quality sequences using Trimmomatic-v0.38 ^43^. Then, the clean Illumina short reads were mapped to the polished contigs using BWA-MEM-v0.7.17 ^44^ with default parameters, based on three iterative rounds of polishing with parameters: ‘--fix all’ by Pilon-v1.23 ^45^ (read length N50 = 41.37 Mb). Finally, the polished contigs were aligned to the Heinz genome to form pseudomolecules, by the software RaGOO-v1.1 ^46^. Embryophyta BUSCO-v5.4.2 (odb10) ^47^ was used to evaluate the integrity of the final assembly.

### PacBio transcriptome processing

The SMRTlink-v6.0.0 pipeline was applied to PacBio *lso-seq* raw data. Firstly, the circular consensus sequence reads (CCSs) were extracted from subreads of BAM files with parameters: minLength= 300, minZScore= -999, minPasses= 1, maxDropFraction= 0.8, minPredictedAccuracy= 0.8, minSnr= 4, noPolish= TURE.

Secondly, CCS reads were divided into full-length non-chimeric (FLNC) reads and non-full-length (nFL) reads, by identifying whether 511 and 311 adapters and the poly(A) tail were present. Then, we obtained consensus isoforms using the ICE algorithm from FLNC reads (Iterative Clustering for Error Correction), which were further polished with nFL reads to get high-quality consensus FLNC reads based on Quiver (parameters: hq_quiver_min_accuracy 0.99, others with default parameters). Finally, the nonredundant FLNC reads were obtained by CD-HIT-v4.8.1 ^48^ with an identity cutoff of 0.95. We also used the software LoRDEC-v0.9 ^49^ to correct mismatch and nucleotide indels in FLNC reads.

### Repetitive DNA identification and Genome annotation

We identified the repetitive elements in the microTom genome based on a combination of *de novo* and homology-based strategies. A *de novo* library was first built through RepeatModeler-v2.0.1 ^50^ and LTR_retriever-v2.9.0 ^51^. The repeat region in the genome was further masked by scanning *de novo* repeat library. Next, we used RepeatMasker-v4.0.9 ^52^ to classify the repeat sequences using the parameters: -poly -html -gff -lib -no_is -xsmall.

The combination of *ab initio*, homology-based and RNA-Seq-based annotation strategy was applied to predict and annotate the protein-coding genes. Firstly, the masked genome was trained by BRAKER2-v2.1.5 ^53^ and GeneMark-ET-v4.69 ^54^ for *ab initio* gene prediction. Secondly, for finding homologous genes, related species like *Solanum tuberosum* (686_v6.1) and *Solanum lycopersicum* Heinz 1706 (691_ITAG4.0), which downloaded from Phytozome v13, were loaded into GenomeThreader-v1.7.1 ^55^ with the parameters: -species arabidopsis -gff3out -intermediate. Thirdly, 81 RNA-Seq (**Supplementary Figure S3 and Table S4**) data sets were used to assist the annotation. These were evaluated by FastQC-v0.11.9 and Trimmomatic-v0.38 ^43^ to trim low-quality and adaptor sequences. Then, these high-quality reads were aligned to the genome by HISAT2-v2.2.1 ^56^. We assembled transcripts with StringTie-v2.1.5 ^57^ and TransDecoder-v5.5.0 (https://github.com/TransDecoder/TransDecoder). The PacBio *lso-seq* was also applied to forecast complete coding sequences (CDS) by PASA-v2.5.2 pipeline ^58^. Finally, the EVidenceModeler-v1.1.1 pipeline was applied to integrate the above-mentioned prediction strategies ^59^. The GTF annotation file, assembled from the Illumina and PacBio data, was used for identifying AS events using the AStalavista v3.2 tool ^60^.

### Functional annotation

Several public databases (PFAM, GO and KEGG) were used to map GO terms for searching functional motifs and domains by InterProScan-v5.50-84.0 ^61^. When the Gene Ontology terms for each gene were obtained, functions of protein-coding genes in microTom genome were annotated by SWISS-PROT database, only retaining the best alignment.

### Identification of SVs and SNPs between reference genomes

Structural variations (SVs) were identified by genome-wide alignment between the microTom and Heinz 1706 (ITAG4.0) genomes using MUMandCo-v3.8 ^62^, which can detect over 1 kb structural variations in length. We also used the MUMmer package-v4.0.0 ^63^ to align these two genomes. The reciprocal best alignments were found using the delta-filter program. Then, the uniquely aligned fragments were used to identify single nucleotide polymorphism (SNP) sites and indels with the show-snp tool.

### Gene expression and construction of co-expression networks

The RNA-seq dataset for gene co-expression analysis was obtained from samples with 22 various tissues/organs and at different developmental stages and/or different treatments. The normalized expression of RNA-seq datawas calculated with FPKM (fragments per kb per million reads) from all samples. FPKM values lower than 1 were filtered out. We used the R package WGCNA (v1.66) ^64^ to construct co-expression networks, and the soft threshold value was calculated by pickSoftThreshold integrated in WGCNA. We used cytoHubba and MCODE to identify candidate hub genes, and used Cytoscape-v3.9.1 ^65^ to visualize the co-expression networks.

### Identification and annotation of Non-coding RNAs

The prediction of protein-coding potential for transcripts generated from Illumina and SMRT data was conducted by integrating CPC2 v0.1, CNCI v2, PLEK v1.2, and PFAM v32 ^66–69^. Candidate lncRNAs were defined as transcripts with the longest representative sequence lacking any open reading frame (ORF) exceeding 100 amino acids, while having a minimum nucleotide sequence length of 200 nts. MiRNAs were identified using miRador (https://github.com/rkweku/miRador) with slight modification ^70^. Specifically, the sequences with the highest abundance of each miRNA locus were added as candidate miRNAs. Candidate miRNAs were annotated as known or novel miRNAs through referencing the latest miRBase version 22.1 ^71^. CIRI2 (v2.0.6), CIRIexplorer2 (v2.4.0), and Find_ciric (v1.2) were combined to detect potential back-splice sites of candidate circRNAs ^72–74^. Reconstruction of full-length circRNAs was achieved using CIRI-full ^75^.

### Identification of targets of miRNAs/lncRNAs

Potential miRNA targets (mRNAs and lncRNAs) were firstly predicted by using the psRNATarget v2 program. Then, the expression values of resultant miRNA-mRNA/lncRNA pairs for six tissues (roots, stems, leaves, flower, immature green fruits, and mature red fruits) were subjected to the spearman correlation coefficient analysis. MiRNA-mRNA/lncRNA pairs were considered to be a candidate interactional miRNA-target when the spearman correlation coefficient was <-0.80 and the p value was <0.01.

To predict lncRNA-mRNA co-expressed pairs, the spearman correlation analysis was performed between lncRNAs and mRNAs in 75 RNA-seq samples. For more accurate prediction of lncRNA-mRNA correlations, we increased the threshold of spearman correlation coefficient| to >0.9, and the p value <0.01. We also screened the results that met the above requirements based on six or more samples with FPKM greater than 1, considering these lncRNA-mRNA pairs to be co-expressed.

## Data availability

The data supporting the findings of this work are available within the paper and its Supplementary information files. The datasets generated and analyzed during this study are available from the corresponding author upon request. All sequencing data generated have been deposited in National Genomics Data Center (https://bigd.big.ac.cn/) with BioProject ID: PRJCA012325 under GSA: CRA008482. The genome assembly, along with the gene models, the functional annotation, and some supplementary information files are available on the FigShare at the link: (https://doi.org/10.6084/m9.figshare.21509817). All these information can also be accessed under GWH: WGS029763 and microTomBase (http://eplant.njau.edu.cn/microTomBase).

## Author contributions

J.Y.X., Z.Q.S., F.C. and Y.V.d.P. conceived the study. Z.Q.S. and Z.Z. collected plant samples. H.Y.F. conducted genome assembly and annotation. H.Y.F., Z.Z., S.Y.H. Y.J.C., and S.X.L conducted analyses of RNA-seq data. Y.H.Z. constructed the database. J.Y.X. and Z.Q.S. drafted the manuscript. All authors read the manuscript and participated in the revision of the manuscript. All authors approved the final manuscript.

## Acknowledgments

This work was supported by grants from the Fundamental Research Funds for the Central Universities (KYCXJC2022003), the National Natural Science Foundation of China (32070243) and the Outstanding Young Teacher of “QingLan Project” of Jiangsu Province. Y.V.d.P. acknowledges funding from the European Research Council (ERC) under the European Union’s Horizon 2020 research and innovation program (No. 833522) and from Ghent University (Methusalem funding, BOF.MET.2021.0005.01). The computational resources and services were provided by the Bioinformatics Center of Nanjing Agricultural University.

## Conflict of interest statement

The authors declare that the research was conducted in the absence of any commercial or financial relationships that could be construed as a potential conflict of interest.

## Supplementary information

Supplementary Fig. S1. Statistics the number of duplicate genes, specific genes and missed genes due to annotation in Heinz and microTom.

Supplementary Fig. S2. The expressed transcripts classified into 123 modules. Supplementary Fig. S3. Information about transcriptome datasets and sequencing strategies.

Supplementary Fig. S4. Evolutionary and divergence time estimation. Phylogeny of eleven species. The numbers on the nodes indicated the divergence time of species (Mya, million years ago), with the confidence range in brackets.

Supplementary Fig. S5. Genomic collinearity analysis between microTom and Heinz. Supplementary Fig. S6. Density of synonymous substitutions per synonymous site (Ks) of collinear paralogous genes.

Supplementary Table S1. Quantitative expression of transcriptome data (Fpkm). Supplementary Table S2. Gene information in the model block.

Supplementary Table S3. A dataset of significantly correlated combinations between non-coding RNAs and their target RNAs.

Supplementary Table S4. Summary of transcriptome mapping for microTom tissues.

## Notes

### Competing Interest Statement

The authors have declared no competing interest.

### Summary of Updates

Updation on microTom database; Remove the data about RNA editing.

## References

1. Klee, H.J. & Giovannoni, J.J. Genetics and control of tomato fruit ripening and quality attributes. Annu Rev Genet 45, 41–59 http://dx.doi.org/10.1146/annurev-genet-110410-132507 (2011).

2. Azzi, L. et al. Fruit growth-related genes in tomato. J Exp Bot 66, 1075-86 http://dx.doi.org/10.1093/jxb/eru527 (2015).

3. Li, Y., et al. MicroTom Metabolic Network: Rewiring Tomato Metabolic Regulatory Network throughout the Growth Cycle. Mol Plant 13, 1203-1218 http://dx.doi.org/10.1016/j.molp.2020.06.005 (2020).

4. Zhu, G. et al. Rewiring of the Fruit Metabolome in Tomato Breeding. Cell 172, 249-261 e12 http://dx.doi.org/10.1016/j.cell.2017.12.019 (2018).

5. Li, J. et al. Biofortified tomatoes provide a new route to vitamin D su fNf iactiPelnacnyts. 8, 611–616 http://dx.doi.org/10.1038/s41477-022-01154-6 (2022).

6. Lozano-Torres, J.L., et al. Dual disease resistance mediated by the immune receptor Cf-2 in tomato requires a common virulence target of a fungus and a nematode. Proc Natl Acad Sci U S A 109, 10119-24 http://dx.doi.org/10.1073/pnas.1202867109 (2012).

7. Weiberg, A. et al. Flungal small RNAs suppress plant immunity by hijacking host RNA interference pathways. Science 342, 118-23 http://dx.doi.org/10.1126/science.1239705 (2013).

8. Chitarra, W. et a lI.nsights on the Impact of Arbuscular Mycorrhizal Symbiosis on Tomato Tolerance to Water Stress. Plant Physiol 171, 1009-23 http://dx.doi.org/10.1104/pp.16.00307 (2016).

9. Liao, D. et a lS. lSPX1-SlPHR complexes mediate the suppression of arbuscular mycorrhizal symbiosis by phosphate repletion in tomato. Plant Cell http://dx.doi.org/10.1093/plcell/koac212 (2022).

10. Alonge, M. et al. Major Impacts of Widespread Structural Variation on Gene Expression and Crop Improvement in Tomato. Cell 182, 145-161 e23 http://dx.doi.org/10.1016/j.cell.2020.05.021 (2020).

11. Gao, L. et al. The tomato pan-genome uncovers new genes and a rare allele regulating fruit flavor. Nat Genet 51, 1044-1051 http://dx.doi.org/10.1038/s41588-019-0410-2 (2019).

12. Zhou, Y. et al. Glraph pangenome captures missing heritability and empowers tomato breeding. Nature 606, 527–534 http://dx.doi.org/10.1038/s41586-022-04808-9 (2022).

13. Wang, W. et al. Genomic variation in 3,010 diverse accessions of Asian cultivated rice. Nature 557, 43–49 http://dx.doi.org/10.1038/s41586-018-0063-9 (2018).

14. Ma, Z. et al. Resequencing a core collection of upland cotton identifies genomic variation and loci influencing fiber quality and yield. Nat Genet 50, 803–813 http://dx.doi.org/10.1038/s41588-018-0119-7 (2018).

15. Song, J.M., et al E. ight high-quality genomes reveal pan-genome architecture and ecotype differentiation of Brassica napus. Nat Plants 6, 34–45 http://dx.doi.org/10.1038/s41477-019-0577-7 (2020).

16. Liu, Y. et al. Pan-Genome of Wild and Cultivated Soybeans. Cell 182, 162-176 e13 http://dx.doi.org/10.1016/j.cell.2020.05.023 (2020).

17. Zeng, Z. et a lT. he RNAome landscape of tomato during arbuscular mycorrhizal symbiosis reveals an evolving RNA layer symbiotic regulatory network. Plant Commun 100429 http://dx.doi.org/10.1016/j.xplc.2022.100429 (2022).

18. Ule, J. & Blencowe, B.J. Alternative Splicing Regulatory Networks: Functions, Mechanisms, and Evolution. Mol Cell76, 329-345 http://dx.doi.org/10.1016/j.molcel.2019.09.017 (2019).

19. Rey, T., et al T. he Medicago truncatula GRAS protein RAD1 supports arbuscular mycorrhiza symbiosis and Phytophthora palmivora susceptibility. Journal of Experimental Botany 68, 5871-5881 http://dx.doi.org/10.1093/jxb/erx398 (2017).

20. Pimprikar, P. et a lA. CCaMK-CYCLOPS-DELLA Complex Activates Transcription of RAM1 to Regulate Arbuscule Branching (vol 26, pg 987, 2016). Curr Biol 26, 1126-112http://dx.doi.org/10.1016/j.cub.2016.04.021 (2016).

21. Xue, L. et a lN. etwork of GRAS Transcription Factors Involved in the Control of Arbuscule Development in Lotus japonicus. Plant Physiology 167, 854-+ http://dx.doi.org/10.1104/pp.114.255430 (2015).

22. Rich, M.K. et al. The Petunia GRAS Transcription Factor ATA/RAM1 Regulates Symbiotic Gene Expression and Fungal Morphogenesis in Arbuscular Mycorrhiza. Plant Physiology 168, 788-+ http://dx.doi.org/10.1104/pp.15.00310 (2015).

23. Gobbato, E., et al. A GRAS-Type Transcription Factor with a Specific Function in Mycorrhizal Signaling. Curr Biol 22, 2236-2241 http://dx.doi.org/10.1016/j.cub.2012.09.044 (2012).

24. Jiang, Y.N. et al. Medicago AP2-Domain Transcription Factor WRI5a Is a Master Regulator of Lipid Biosynthesis and Transfer during Mycorrhizal Symbiosis. Molecular Plant 11, 1344-1359 http://dx.doi.org/10.1016/j.molp.2018.09.006 (2018).

25. Park, J.W., Yang, J. & Xu, R.H. Paired Box Protein 6 Alternative Splicing and Corneal Development. Stem Cells Dev 27, 367-377 http://dx.doi.org/10.1089/scd.2017.0283 (2018).

26. Baralle, F.E. & Giudice, J. Alternative splicing as a regulator of development and tissue identity. Nat Rev Mol Cell Bio 18, 437-451 http://dx.doi.org/10.1038/nrm.2017.27 (2017).

27. Zhang, Y.J., Qian, J.J., Gu, C.Y. & Yang, Y. Alternative splicing and cancer: a systematic review. Signal Transduct Tar 6, http://dx.doi.org/ARTN 78 10.1038/s41392-021-00486-7 (2021).

28. Lukas, J., et al. Alternative and aberrant messenger RNA splicing of the mdm2 oncogene in invasive breast cancer. Cancer Res 61, 3212–3219 <Go to ISI>://WOS:000167967100066 (2001).

29. Laloum, T., Martin, G. & Duque, P. Alternative Splicing Control of Abiotic Stress Responses. Trends Plant Sci 23, 140-150 http://dx.doi.org/10.1016/j.tplants.2017.09.019 (2018).

30. John, S., Olas, J.J. & Mueller-Roeber, B. Regulation of alternative splicing in response to temperature variation in plants. Journal of Experimental Botany 72, 6150-6163 http://dx.doi.org/10.1093/jxb/erab232 (2021).

31. Zhang, X.C. & Gassmann, W. Alternative splicing and mRNA levels of the disease resistance gene RPS4 are induced during Defense responses(1[W][OA]). Plant Physiology 145, 1577-1587 http://dx.doi.org/10.1104/pp.107.108720 (2007).

32. Song, X., Li, Y., Cao, X. & Qi, Y. MicroRNAs and Their Regulatory Roles in Plant-Environment Interactions. Annu Rev Plant Biol 70, 489–525 http://dx.doi.org/10.1146/annurev-arplant-050718-100334 (2019).

33. Yu, Y., Zhang, Y., Chen, X. & Chen, Y. Plant Noncoding RNAs: Hidden Players in Development and Stress Responses. A n n u R e v http://dx.doi.org/10.1146/annurev-cellbio-100818-125218<x> (2019). 35,C407-4e31 l l D e v

34. Dong, F., et al. Differential expression of microRNAs in tomato leaves treated with different light qualities. BMC Genomics 21, 37 http://dx.doi.org/10.1186/s12864-019-6440-4 (2020).

35. Cao, H. et al. miRNA expression profiling and zeatin dynamic changes in a new model system of in vivo indirect regeneration of tomato. PLoS One 15, e0237690 http://dx.doi.org/10.1371/journal.pone.0237690 (2020).

36. Feng, J., Liu, S., Wang, M., Lang, Q. & Jin, C. Identification of microRNAs and their targets in tomato infected with Cucumber mosaic virus based on deep sequencing. Planta 240, 1335-52 http://dx.doi.org/10.1007/s00425-014-2158-3 (2014).

37. Ferreira e Silva, G.F., et al. microRNA156-targeted SPL/SBP box transcription factors regulate tomato ovary and fruit development. Plant J 78, 604-18 http://dx.doi.org/10.1111/tpj.12493 (2014).

38. da Silva, E.M., et al m. icroRNA159-targeted SlGAMYB transcription factors are required for fruit set in tomato. Plant J 92, 95-109 http://dx.doi.org/10.1111/tpj.13637 (2017).

39. Lamin-Samu, A.T., Zhuo, S., Ali, M. & Lu, G. Long non-coding RNA transcriptome landscape of anthers at different developmental stages in response to drought stress in tomato. Genomics 114, 110383 http://dx.doi.org/10.1016/j.ygeno.2022.110383 (2022).

40. Fabian, M.R., Sonenberg, N. & Filipowicz, W. Regulation of mRNA translation and stability by microRNAs. A n n u http://dx.doi.org/10.1146/annurev-biochem-060308-103103<x> (2010). 79, 351-R79 e v

41. Li, H. New strategies to improve minimap2 alignment accuracy. Bioinformatics 37, 4572-4574 http://dx.doi.org/10.1093/bioinformatics/btab705 (2021).

42. Hu, J., Fan, J., Sun, Z. & Liu, S. NextPolish: a fast and efficient genome polishing tool for long-read assembly. Bioinformatics 36, 2253-2255 http://dx.doi.org/10.1093/bioinformatics/btz891 (2019).

43. Bolger, A.M., Lohse, M. & Usadel, B. Trimmomatic: a flexible trimmer for Illumina sequence data. Bioinformatics 30, 2114-20 http://dx.doi.org/10.1093/bioinformatics/btu170 (2014).

44. Li, H. & Durbin, R. Fast and accurate short read alignment with Burrows-Wheeler transform. Bioinformatics 25, 1754-60 http://dx.doi.org/10.1093/bioinformatics/btp324 (2009).

45. Walker, B.J. et al. Pilon: an integrated tool for comprehensive microbial variant detection and genome assembly improvement. PLoS One. Preprint at http://europepmc.org/abstract/MED/25409509 https://doi.org/10.1371/journal.pone.0112963 https://europepmc.org/articles/PMC4237348 https://europepmc.org/articles/PMC4237348?pdf=render (2014).

46. Alonge, M., et alR. aGOO: fast and accurate reference-guided scaffolding of draft genomes. Genome Biology20, 224 http://dx.doi.org/10.1186/s13059-019-1829-6 (2019).

47. Manni, M., Berkeley, M.R., Seppey, M. & Zdobnov, E.M. BUSCO: Assessing Genomic Data Quality and Beyond. Curr Protoc 1, e323 http://dx.doi.org/10.1002/cpz1.323 (2021).

48. Fu, L., Niu, B., Zhu, Z., Wu, S. & Li, W. CD-HIT: accelerated for clustering the next-generation sequencing data. Bioinformatics 28, 3150-3152 http://dx.doi.org/10.1093/bioinformatics/bts565 (2012).

49. Salmela, L. & Rivals, E. LoRDEC: accurate and efficient long read error correction. BIOINFORMATICS 30, 3506-3514 http://dx.doi.org/10.1093/bioinformatics/btu538 (2014).

50. Bao, Z. & Eddy, S.R. Automated de novo identification of repeat sequence families in sequenced genomes. Genome research 12, 1269-1276 http://dx.doi.org/10.1101/gr.88502 (2002).

51. Ou, S. & Jiang, N. LTR_retriever: A Highly Accurate and Sensitive Program for Identification of Long Terminal Repeat Retrotransposons. Plant Physiol 176, 1410-1422 http://dx.doi.org/10.1104/pp.17.01310 (2018).

52. Tarailo-Graovac, M. & Chen, N. Using RepeatMasker to identify repetitive elements in genomic sequences. Curr Protoc Bioinformatics Chapter 4, Unit 4.10 http://dx.doi.org/10.1002/0471250953.bi0410s25 (2009).

53. Brůna, T., Hoff, K.J., Lomsadze, A., Stanke, M. & Borodovsky, M. BRAKER2: automatic eukaryotic genome annotation with GeneMark-EP+ and AUGUSTUS supported by a protein database. NAR Genomics and Bioinformatics 3, http://dx.doi.org/10.1093/nargab/lqaa108 (2021).

54. Lomsadze, A., Burns, P.D. & Borodovsky, M. Integration of mapped RNA-Seq reads into automatic training of eukaryotic gene finding algorithm. Nucleic Acids Res 42, e119 http://dx.doi.org/10.1093/nar/gku557 (2014).

55. Gremme, G., Brendel, V., Sparks, M.E. & Kurtz, S. Engineering a software tool for gene structure prediction in higher organisms. Information and Software Technology 47, 965-978 http://dx.doi.org/https://doi.org/10.1016/j.infsof.2005.09.005 (2005).

56. Kim, D., Paggi, J.M., Park, C., Bennett, C. & Salzberg, S.L. Graph-based genome alignment and genotyping with HISAT2 and HISAT-genotype. Nat Biotechnol 37, 907–915 http://dx.doi.org/10.1038/s41587-019-0201-4 (2019).

57. Pertea, M. et al. StringTie enables improved reconstruction of a transcriptome from RNA-seq reads. NATURE BIOTECHNOLOGY33, 290-+ http://dx.doi.org/10.1038/nbt.3122 (2015).

58. Haas, B.J. et a lIm. proving the Arabidopsis genome annotation using maximal transcript alignment assemblies. NUCLEIC ACIDS RESEARCH 31, 5654-5666 http://dx.doi.org/10.1093/nar/gkg770 (2003).

59. Haas, B.J. et al. Automated eukaryotic gene structure annotation using EVidenceModeler and the program to assemble spliced alignments. GENOME BIOLOGY (Article) 9, R7 http://dx.doi.org/10.1186/gb-2008-9-1-r7 (2008).

60. Foissac, S. & Sammeth, M. ASTALAVISTA: dynamic and flexible analysis of alternative splicing events in custom gene datasets. Nucleic Acids Res 35, W297-9 http://dx.doi.org/10.1093/nar/gkm311 (2007).

61. Zdobnov, E.M. & Apweiler, R. InterProScan - an integration platform for the signature-recognition methods in InterPro. BIOINFORMATICS 17, 847-848 http://dx.doi.org/10.1093/bioinformatics/17.9.847 (2001).

62. O’Donnell, S. & Fischer, G. MUM<Co: accurate detection of all SV types through whole-genome alignment. Bioinformatics 36, 3242-3243 http://dx.doi.org/10.1093/bioinformatics/btaa115 (2020).

63. Kurtz, S. et al. Versatile and open software for comparing large genomes. GENOME BIOLOGY 5, R12 http://dx.doi.org/10.1186/gb-2004-5-2-r12 (2004).

64. Langfelder, P. & Horvath, S. WGCNA: an R package for weighted correlation network analysis. BMC BIOINFORMATICS 9, 559 http://dx.doi.org/10.1186/1471-2105-9-559 (2008).

65. Shannon, P. et al. Cytoscape: A software environment for integrated models of biomolecular interaction networks. Genome Research 13, 2498-2504 http://dx.doi.org/10.1101/gr.1239303 (2003).

66. Li, A., Zhang, J. & Zhou, Z. PLEK: a tool for predicting long non-coding RNAs and messenger R N A s b a s e d o n a n i mBMpC Brioinoforvmateicsd http://dx.doi.org/10.1186/1471-2105-15-311<x> (2014). 15k, 3-11 m e r s c h e m

67. Kong, L. et al. CPC: assess the protein-coding potential of transcripts using sequence features and support vector machine. Nucleic Acids Res 35, W345–9 http://dx.doi.org/10.1093/nar/gkm391 (2007).

68. Finn, R.D. et al. Pfam : the proteinfam iNluci leeicsAcidds aRets a b a42s, De2.22-30 http://dx.doi.org/10.1093/nar/gkt1223 (2014).

69. Sun, L. et a lU.tilizing sequence intrinsic composition to classify protein-coding and long non-coding transcripts. Nucleic Acids Res 41, e166 http://dx.doi.org/10.1093/nar/gkt646 (2013).

70. Hammond, R.K., Gupta, P., Patel, P. & Meyers, B.C. miRador: a fast and precise tool for the prediction of plant miRNAs. bioRxiv http://dx.doi.org/doi : https://doi.org/10.1101/2021.03.24.436803 (2021).

71. Kozomara, A., Birgaoanu, M. & Griffiths-Jones, S. miRBase: from microRNA sequences to function. Nucleic Acids Res 47, D155–D162 http://dx.doi.org/10.1093/nar/gky1141 (2019).

72. Memczak, S., et al. Circular RNAs are a large class of animal RNAs with regulatory potency. Nature 495, 333–8 http://dx.doi.org/10.1038/nature11928 (2013).

73. Gao, Y., Zhang, J. & Zhao, F. Circular RNA identification based on multiple seed matching. Brief Bioinform 19, 803–810 http://dx.doi.org/10.1093/bib/bbx014 (2018).

74. Zhang, X.O. et al. Diverse alternative back-splicing and alternative splicing landscape of circular RNAs. Genome Res 26, 1277-87 http://dx.doi.org/10.1101/gr.202895.115 (2016).

75. Zheng, Y., Ji, P., Chen, S., Hou, L. & Zhao, F. Reconstruction of full-length circular RNAs enables isoform-level quantification. Genome Med 11, 2 http://dx.doi.org/10.1186/s13073-019-0614-1 (2019).

