## Supplementary material for "Comprehensive regulatory networks for tomato organ development based on the genome and RNAome of microTom tomato": sulp.fig.docx

**Supplementary Fig. S1. Statistics the number of duplicate genes, specific genes and missed genes due to annotation in Heinz and microTom.**

**Supplementary Fig. S2. The expressed transcripts classified into 123 modules.**

**Supplementary Fig. S3. Information about transcriptome datasets and sequencing strategies.**

**Supplementary Fig. S4. Evolutionary and divergence time estimation. Phylogeny of eleven species. The numbers on the nodes indicated the divergence time of species (Mya, million years ago), with the confidence range in brackets.**

**Supplementary Fig. S5. Genomic collinearity analysis between microTom and Heinz.**

**Supplementary Fig. S6. Density of synonymous substitutions per synonymous site (Ks) of collinear paralogous genes.**

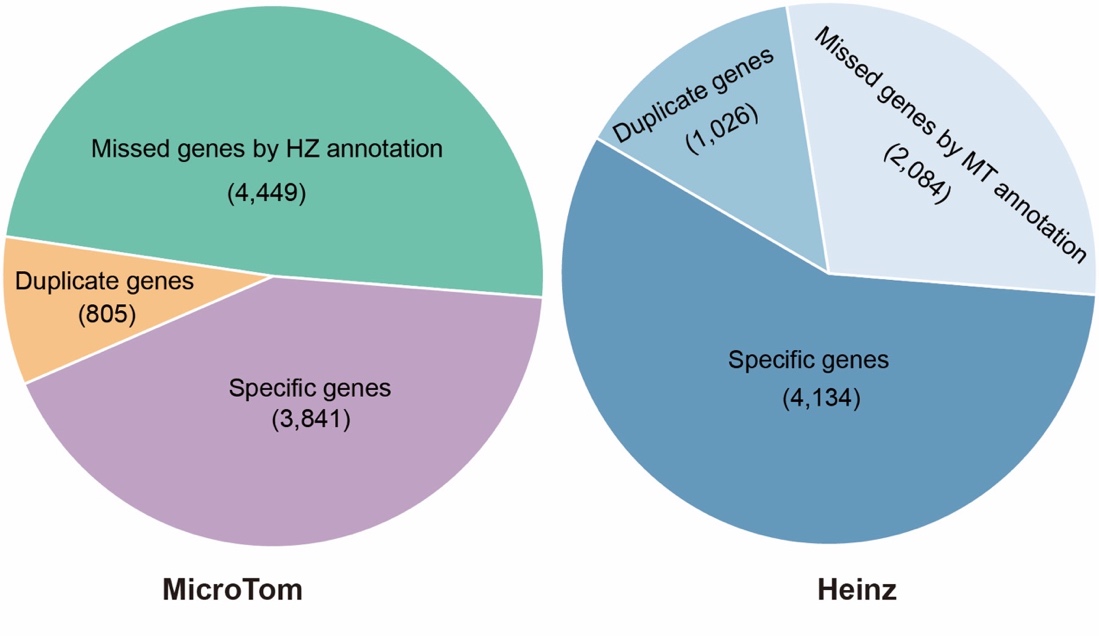

**Supplementary Fig. S1. Statistics the number of duplicate genes, specific genes and missed genes due to annotation in Heinz and microTom.**

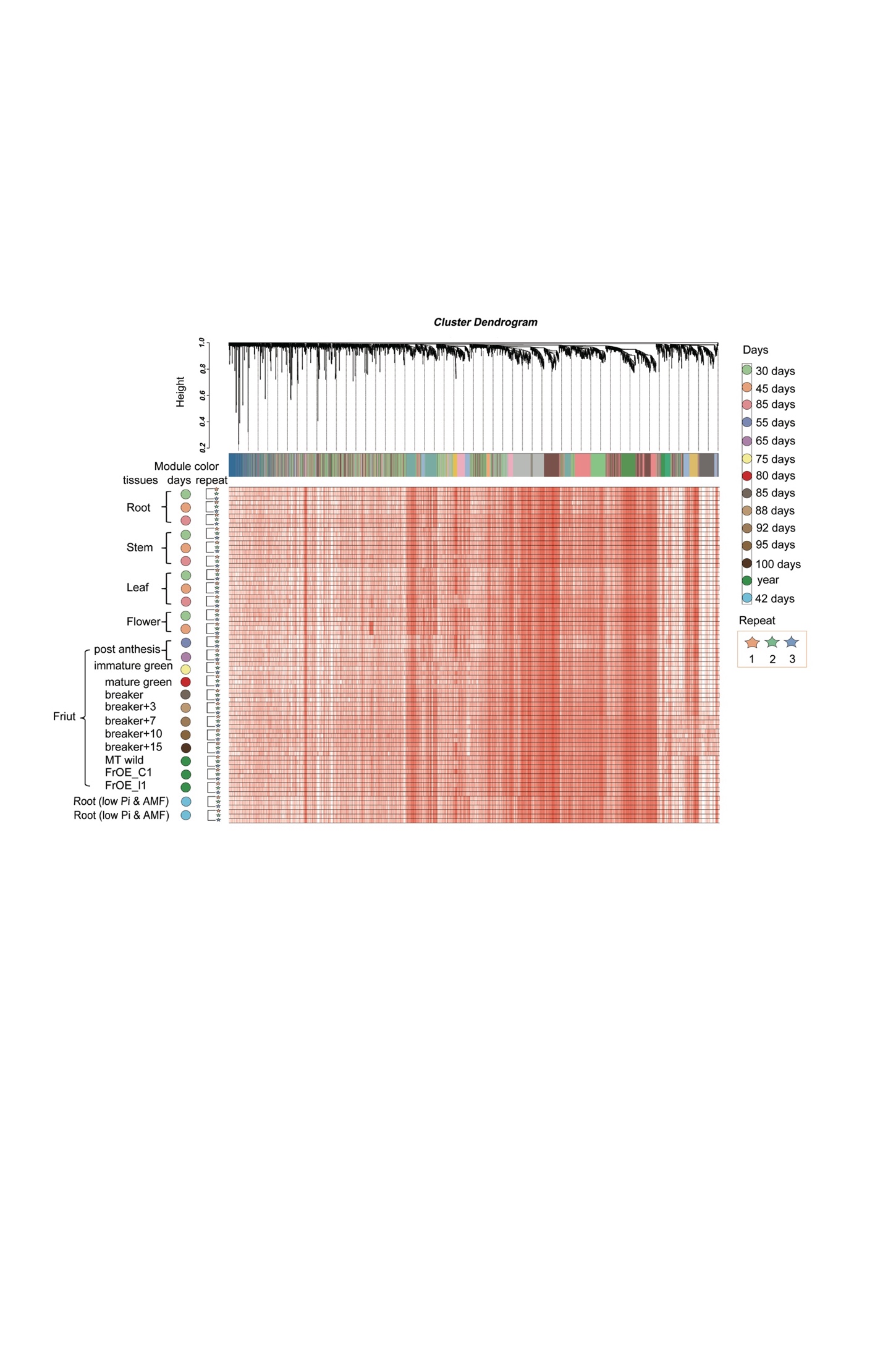

**Supplementary Fig. S2. The expressed transcripts classified into 123 modules.**

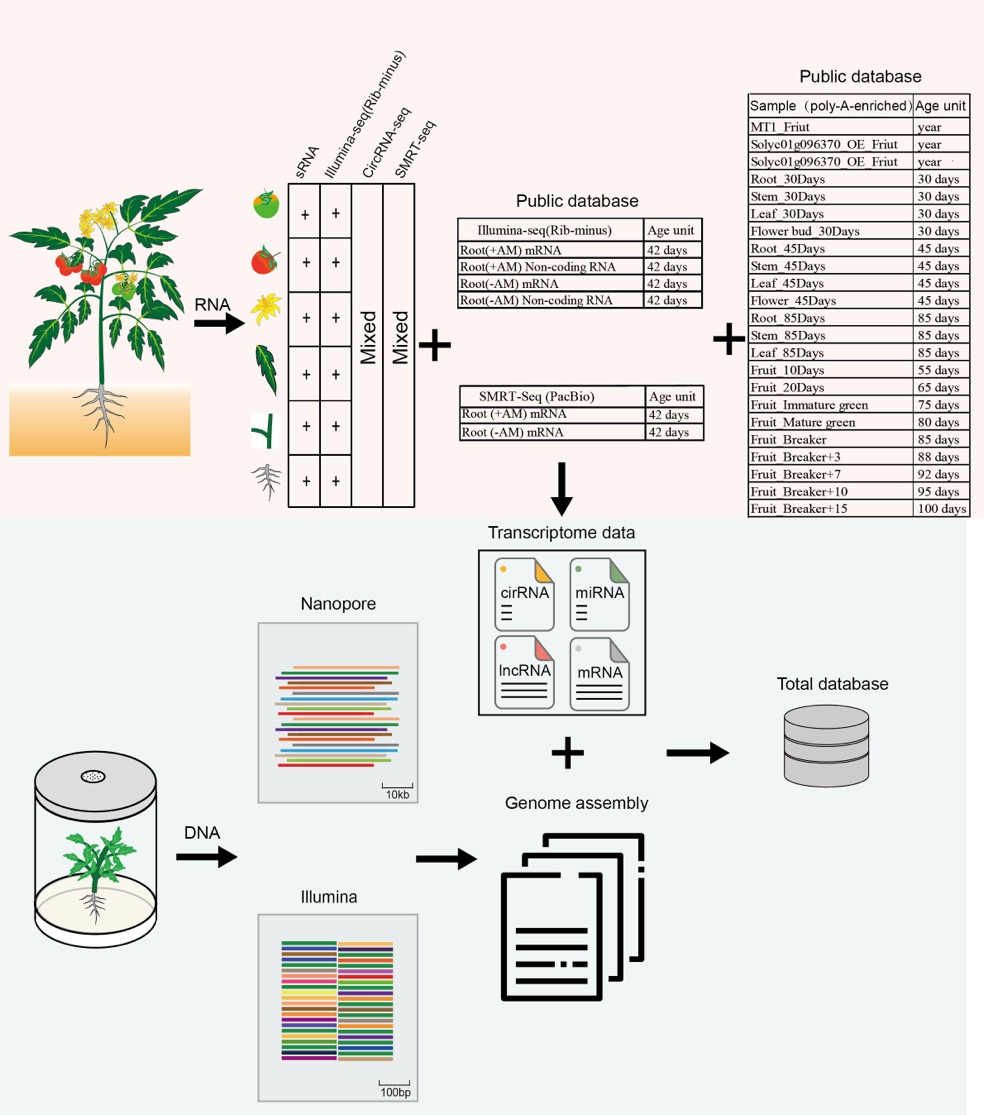

**Supplementary Fig. S3. Information about transcriptome datasets and sequencing strategies.**

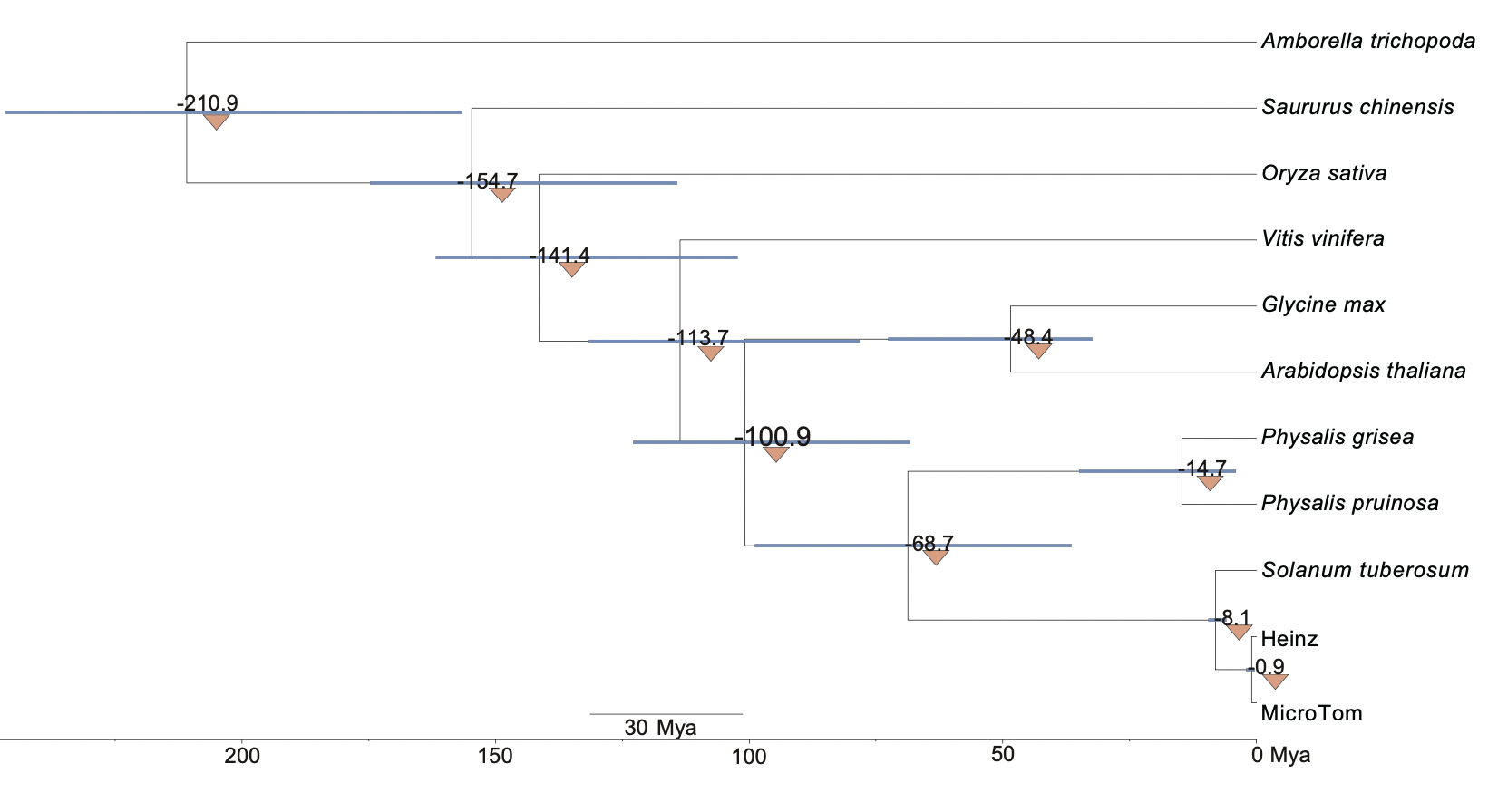

**Supplementary Fig. S4. Evolutionary and divergence time estimation. Phylogeny of eleven species. The numbers on the nodes indicated the divergence time of species (Mya, million years ago), with the confidence range in brackets.**

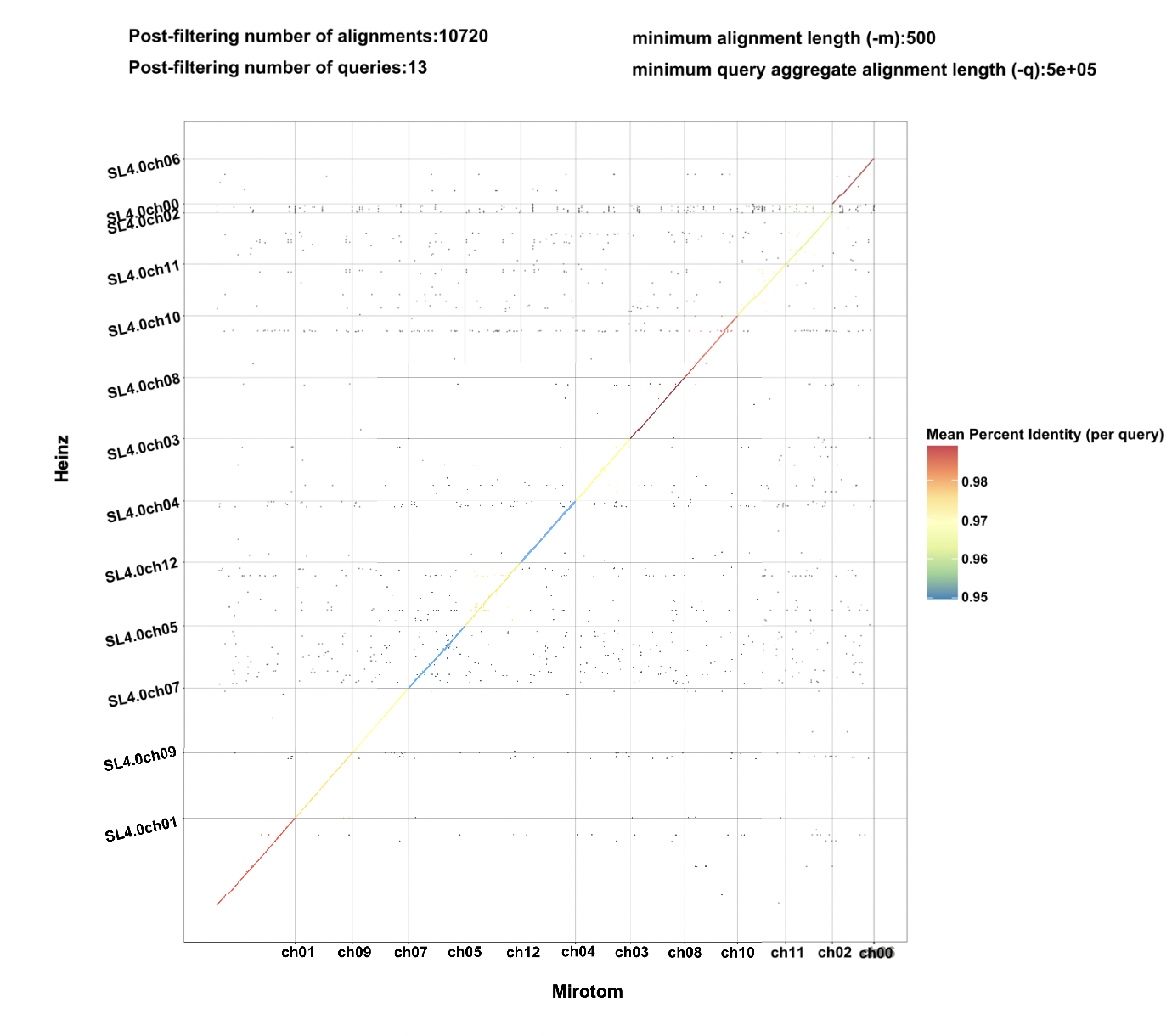

**Supplementary Fig. S5. Genomic collinearity analysis between microTom and Heinz.**

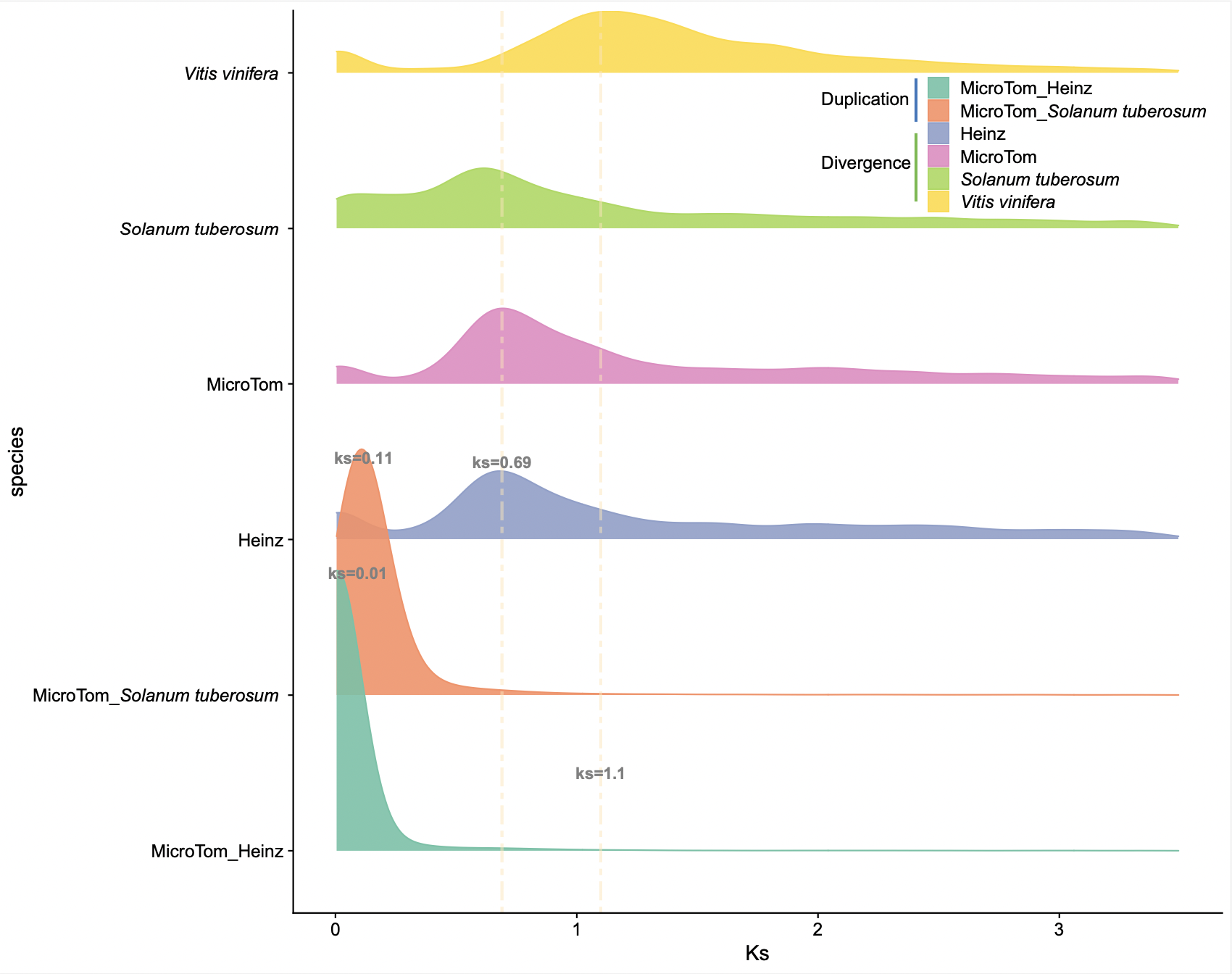

**Supplementary Fig. S6. Density of synonymous substitutions per synonymous site (Ks) of collinear paralogous genes.**
